# Light-based juxtacrine signaling between synthetic cells

**DOI:** 10.1101/2024.01.05.574425

**Authors:** Hossein Moghimianavval, Kyle J. Loi, Sung-Won Hwang, Yashar Bashirzadeh, Allen P. Liu

## Abstract

Cell signaling through direct physical cell-cell contacts plays vital roles in biology during development, angiogenesis, and immune response. Intercellular communication mechanisms between synthetic cells constructed from the bottom up are majorly reliant on diffusible chemical signals, thus limiting the range of responses in receiver cells. Engineering contact-dependent signaling between synthetic cells promises to unlock more complicated signaling schemes with different types of responses. Here, we design and demonstrate a light-activated contact-dependent communication tool for synthetic cells. We utilize a split bioluminescent protein to limit signal generation exclusively to contact interfaces of synthetic cells, driving the recruitment of a photoswitchable protein in receiver cells, akin to juxtacrine signaling in living cells. Our modular design not only demonstrates contact-dependent communication between synthetic cells but also provides a platform for engineering orthogonal contact-dependent signaling mechanisms.

## Introduction

Intercellular communication is a key characteristic of multicellular life. Cells utilize various intercellular communication mechanisms for exchanging information regarding their state, the density of their cellular community, and the presence of pathogens or competing cells^1,2^. While communication through diffusible chemicals is the primary means of signaling in prokaryotes^3^, higher-level organisms like metazoans have amassed a wide range of sophisticated communication mechanisms for short- and long-distance signaling^4^. Specifically, contact-dependent or juxtacrine communication has emerged during evolution as a mechanism for highly specific and targeted communication, especially when signaling results in dramatic responses^5,6^. Notch-Delta signaling is an example of contact-dependent communication with essential roles in gene regulation during development^7,8^. In Notch signaling, direct cell-cell contact enables protein-protein interaction at the contact interface of two cells. Upon interaction between the extracellular domains of Notch and Delta, the intracellular domain of Notch in the receiving cell is proteolytically released which functions to regulate transcription. Similar to Notch signaling, other processes such as T-cell activation^9,10^ and signaling through tight junctions^11^ and gap junctions^12^ rely on cell-cell interface formation and juxtaposed protein-protein interactions for signaling.

Reconstitution of intercellular communication in synthetic cells has recently gained increasing interest^13–15^. The development of synthetic cells with functionalities ranging from light-driven energy generation^16^ to biocomputing^17,18^ and biosensing^19^ has propelled the capabilities of synthetic cells in various applications such as drug delivery and energy regeneration. Therefore, many recent endeavors have focused on implementing communication mechanisms in synthetic cells to enable engineering smart synthetic multicellular systems and synthetic cell-natural cell communication for therapeutic purposes^20^. While initial development in synthetic cell communication utilized the release of membrane-permeable chemical inducers from sender cells to regulate gene expression in receiver cells^21–25^, more sophisticated mechanisms relying on membrane pores^26–28^, CRISPR-Cas systems^29^, or signal amplification^30^ were later developed. Further, the inclusion of light-sensitive elements in the signaling cascade has enabled light-based communication among both communities of synthetic cells as well as synthetic and natural cells^31–36^.

A few synthetic cell contact-dependent communication systems have been engineered previously^26,37–39^, although mostly in a network of connected aqueous droplets in oil. Such studies have relied on membrane pores such as α-hemolysin for chemical communication between adjacent synthetic cells. This has effectively limited the communication between synthetic cells to chemical signaling. The development of SynNotch in natural cells opened up an avenue for engineering cellular communities based on contact-dependent communication for different purposes^40^. However, a similar contact-dependent signaling scheme for synthetic cells that utilizes protein-protein interactions at the contact interface between synthetic cells is yet to be realized.

Here, we engineer a light-based contact-dependent communication system for synthetic cells. We utilized our previously developed strategy^41^ for membrane-membrane interface functionalization and repurposed it to engineer a modular contact-dependent communication system based on intrinsically generated light as a signal. We leveraged optogenetic tools to link the signal, light, to protein recruitment to the membrane interface, supported by mathematical modeling, imitating juxtacrine signaling. The modularity of the design allows for orthogonal contact-dependent signaling schemes, thus paving the way for engineering complex communication pathways between synthetic cells and between synthetic and natural cells in the future.

## Results

### Designing a contact-dependent light-based communication mechanism between synthetic cells

Protein modules that send and receive light are crucial building blocks of a light-based signaling mechanism. In addition, for the signaling cascade to be contact-dependent, such protein modules need to be activated and trigger signaling only when synthetic cells are in contact with each other (**Fig. 1** as described below). We utilized giant unilamellar vesicles (GUVs) as model synthetic cells and sought out appropriate proteins with certain properties that can satisfy all aforementioned conditions.

**Figure 1.**
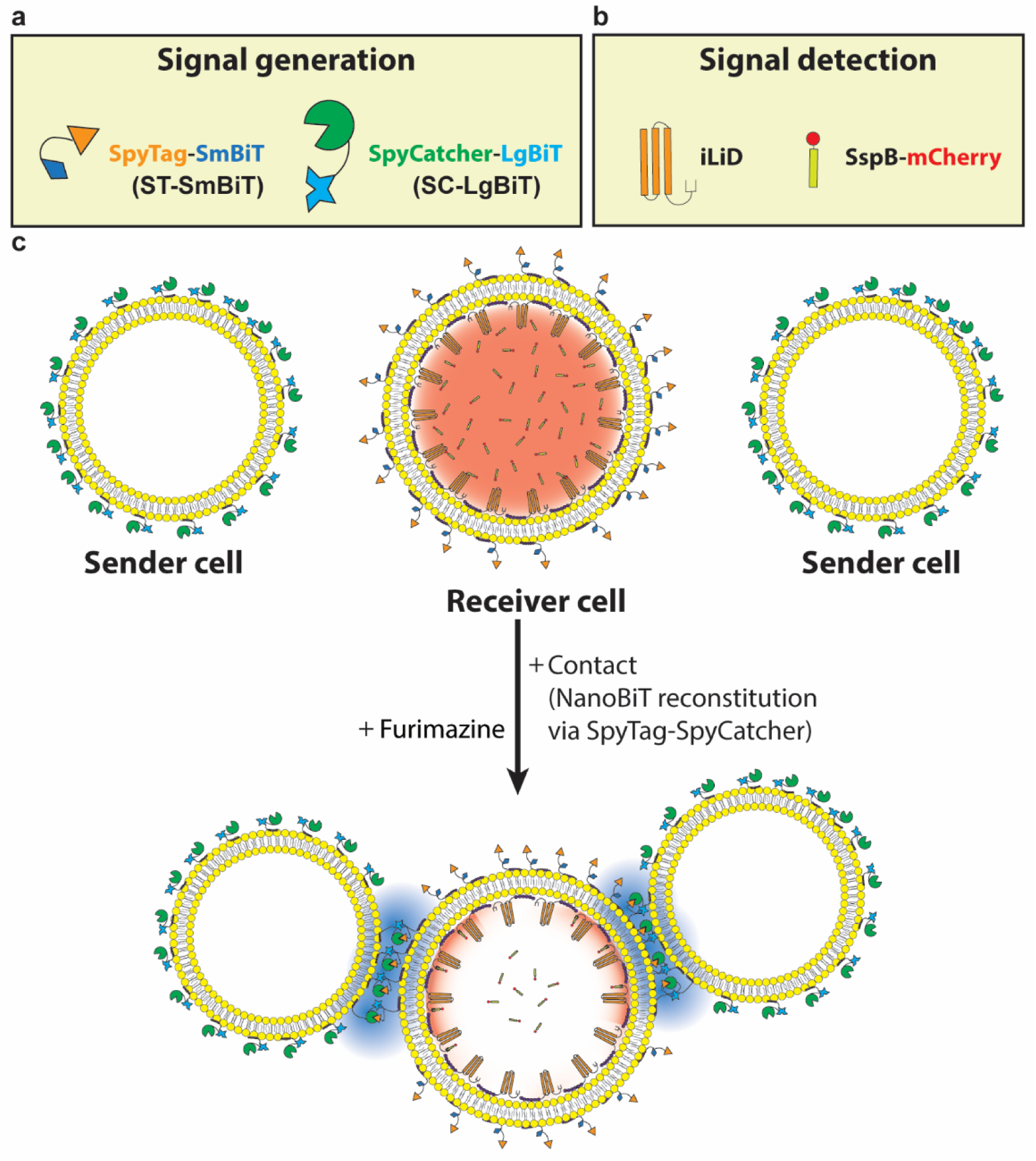
Schematic representation of light-based synthetic cell communication at synthetic cell membrane interfaces. **a,** Signal generation module consists of ST-SmBiT and SC-LgBiT proteins. SpyTag-SpyCatcher interaction brings the SmBiT and LgBiT fragments close to each other, thereby promoting reconstitution of functional NanoBiT luciferase. **b,** Signal detection relies on light-activation of iLID which induces its dimerization with its binding partner, SspB. SspB is translationally fused to a fluorescent mCherry protein which allows detection of SspB translocation upon signal reception. **c,** Direct cell-cell contact between sender and receiver cells causes SpyTag-SpyCatcher interaction, thus signal generation in the presence of furimazine which in turn, promotes iLID activation and translocation of SspB-mCherry from the lumen of receiver cells to the membrane.

Bioluminescent proteins such as *Nanoluc* (NLuc), *Gaussia* (GLuc), and *Renilla* (RLuc) luciferases are ideal candidates for generating blue light through chemiluminescence reactions. The blue light emitted by these proteins can then activate photosensitive elements such as light-oxygen-voltage sensing (LOV) domains that are present in a variety of different photoactivatable proteins widely used for optogenetics applications^42^. Recently, light-emitting synthetic cells encapsulating GLuc that could activate soil fungi *Trichoderma atroviride* were engineered by Adir *et al.*^35^. Additionally, Adir *et al*. demonstrated photoactivation of the light-sensitive transcriptional element EL222 by GLuc bioluminescence inside synthetic cells^35^. Thus, the previous successful utilization of luciferases in synthetic cell development made them a prime choice as signal-generating molecules. Among bioluminescent proteins, NLuc stood out due to its brightness and higher stability compared to its counterparts^43^. NLuc catalyzes the oxidation of its substrate, furimazine, in an ATP-independent manner which results in blue light emission at 450 nm.

Additionally, we wanted the signal generation to only occur when the synthetic cells are in contact with each other. While NLuc had desirable bioluminescence properties for our design, its function would not be restricted to synthetic cell-cell contact interfaces. Inspired by our previous work on developing InterSpy^41^ as a strategy for programmable activation of a split fluorescent protein in membrane-membrane interfaces, we reasoned that a split bioluminescent protein that is activated only at synthetic cell-cell interfaces would be perfectly suited for our design requirements.

A split version of NLuc called Nanoluc Binary Technology (NanoBiT) was designed for reporting protein-protein interactions by Dixon *et al*^44^. NanoBiT consists of a large protein composed of the first 9 beta-sheets of NLuc named large bit (LgBiT) and a small peptide called small bit (SmBiT) that contains NLuc’s 10^th^ beta-sheet strand. LgBiT and SmBiT are designed such that they do not have inherent bioluminescent activity in the presence of furimazine and have low affinity for each other. However, LgBiT and SmBiT complementation and bioluminescence activation occur when the two fragments are brought together via protein-protein interactions. We rationalized that by artificially creating some form of protein-protein interactions between the NanoBiT components exclusively at membrane-membrane interfaces, bioluminescence activity can be reconstituted in synthetic cell-cell contact interfaces. Since InterSpy relies on SpyTag-SpyCatcher interaction for complementation of a split fluorescent cherry protein (sfCherry) in membrane-membrane interfaces^41^, it stood out as an ideal tool for creating the protein-protein interaction required for NanoBiT reconstitution at contact interfaces between sender and receiver synthetic cells. We hypothesized that replacing small and large fragments of sfCherry with SmBiT and LgBiT, respectively, will lead to NanoBiT reconstitution through SpyTag-SpyCatcher interaction exclusively at synthetic cell-cell interfaces. We called the protein made by the fusion of SpyTag and SmBiT as ST-SmBiT and SpyCatcher and LgBiT as SC-LgBiT (**Fig. 1a**).

For signal reception, we selected an improved light-induced dimerizing protein (iLID) for its compatibility with NLuc bioluminescence wavelength, high sensitivity, rapid kinetics, and high affinity for its binding partner, SspB^45^. iLID is a relatively small protein that is made from the LOV2 domain from *Avena sativa* (AsLOV2) with seven residues of *Escherichia coli* SsrA peptide at its C-terminus. The SsrA strand is sterically inaccessible to its dimerizing partner SspB in the dark. However, upon irradiation of blue light, the AsLOV2 domain undergoes structural rearrangement that releases the SsrA peptide, leading to iLID-SspB dimerization^45^. Chakraborty *et al*. used iLID and SspB Nano as adherent molecules between synthetic cell communities to demonstrate light-activated communication between synthetic cells^32,36^. In their work, one group of synthetic cells encapsulated RLuc to emit light, thereby activating iLID-SspB dimerization and adhesion between synthetic cells. Similarly, iLID-SspB Nano binding on the outer membrane of synthetic cells was demonstrated by Adir *et al.*^35^ where N-terminal fusion of GLuc to iLID made iLID activation through GLuc bioluminescence possible. These studies provided a strong motivation to exploit iLID as the signal receptor in our design (**Fig. 1b**).

Lastly, we designed the light-based signaling system such that it resulted in a change in receiver synthetic cells that can be visualized using fluorescence microscopy. In our design, receiver synthetic cells encapsulated membrane-bound iLID and cytosolic SspB Nano-mCherry (referred to as SspB-mCherry hereafter) while the outer membrane of sender and receiver cells were decorated with SC-LgBiT and ST-SmBiT, respectively. Therefore, when a receiver synthetic cell forms a contact interface with a sender synthetic cell, the NanoBiT activation and bioluminescence through SpyTag-SpyCatcher interaction drives iLID-SspB dimerization, leading to membrane-recruitment of mCherry (**Fig. 1c**) which can be detected via fluorescence microscopy.

### SpyTag-SpyCatcher-mediated NanoBiT reconstitution

To test our hypothesis on whether SpyTag-SpyCatcher interaction can lead to functional NanoBiT reconstitution, we made constructs SC-LgBiT and ST-SmBiT by fusing SpyCatcher and SpyTag to the N-terminus of LgBiT and C-terminus of SmBiT, respectively. Since in our design, SC-LgBiT and ST-SmBiT were eventually on the outer surface of sender and receiver synthetic cells (**Fig. 1c**), we designed the constructs such that SC-LgBiT and ST-SmBiT had a C-terminal and a N-terminal 6xHis tag, respectively, so that the molecules could bind to lipids with NTA-Ni headgroups. We then asked whether ST-SmBiT, SC-LgBiT, a 1:1 mixture of ST-SmBiT and SC-LgBiT, or a control experiment where SC-LgBiT was mixed with SmBiT, lacking the SpyTag domain, have bioluminescence activity. To detect bioluminescence, we utilized Nano-Glo assay (**Fig. 2a**). After we made mixtures of 500 nM of the different proteins or their combination and incubated them to ensure SpyTag-SpyCatcher bond formation, we observed that only the mixture of ST-SmBiT and SC-LgBiT demonstrated bioluminescence for about half an hour once furimazine was added (**Fig. 2b**). Notably, the mixture with SC-LgBiT and SmBiT did not show any bioluminescence signal, highlighting the low affinity of SmBiT and LgBiT and the requirement of SpyTag-SpyCatcher isopeptide bond formation in reconstituting NanoBiT as a functional luciferase.

**Figure 2.**
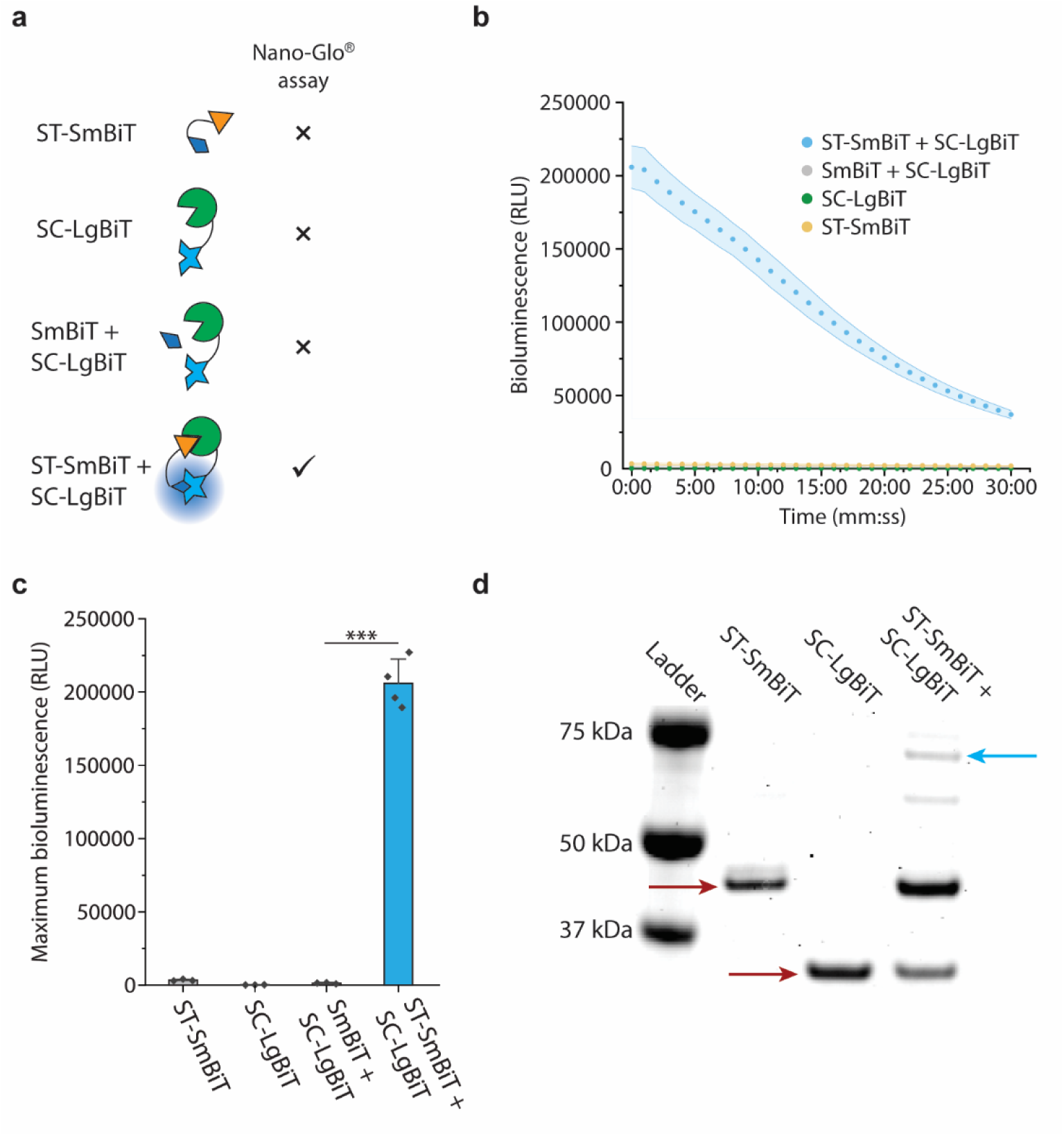
Reconstitution of NanoBIT activity using SpyTag and SpyCatcher. **a**, Schematic of the luciferase assay based on Promega Nano-Glo assay. The left column illustrates different combinations of proteins while the right column represents expected readouts from Nano-Glo assay. Functional reconstitution of NanoBiT only occurs in the presence of protein-protein interaction induced by SpyTag-SpyCatcher dimerization. **b,** The bioluminescence readout from 4 different reactions represented in **a**. Data is represented as mean ± S.D., *n* ≥ 3. **c,** Bar plots representing the maximum bioluminescence signal from four different reactions illustrated in **a**. The error bar represents the standard deviation, n ≥ 3. **d,** In-gel fluorescence imaging of the Coomassie-stained ladder (lane 1), ST-SmBiT (lane 2, red arrow), SC-LgBiT (lane 3, red arrow), and the mixture of SC-LgBiT and ST-SmBiT forming a dimer (lane 4, cyan arrow). *p*-value is calculated using a two-tailed Welch’s *t*-test. *** represents *p* < 0.001.

The highest bioluminescence was detected from the reaction containing the 1:1 mixture of SC-LgBiT and ST-SmBiT while the other conditions demonstrated virtually no bioluminescence activity (**Fig. 2c**). Using bioluminescence imaging, we measured the total photon flux from bioluminescence reactions catalyzed by NLuc or reconstituted NanoBiT to compare light emittance of NanoBiT with its full-length counterpart, NLuc (**Figs. S1a and S1b**). We observed that NLuc consistently had higher maximum total photon flux over a range of different concentrations (**Fig. S1c**). This is expected as Dixon *et al*. also showed that lysates of cells transfected with equal amounts of NanoBiT DNA in the presence of rapamycin demonstrated lower bioluminescence activity compared to lysates with NLuc ^44^. We also attempted to compare the kinetics of NLuc and NanoBiT catalytic reactions. We correlated the measured total photon flux to the exponential decay of the substrate through its exhaustion by NLuc or NanoBiT (**Fig. S2a**). By modeling the bioluminescence reaction via this approach, we calculated the catalytic rate constants of NLuc and NanoBiT. We found that NLuc showed a higher rate constant than NanoBiT (∼38,207 ± 2,450 M^-1^s^-1^ vs. ∼10,644 ± 1,734 M^-1^s^-1^ (95% confidence interval)) (**Figs. S2b and S2c**).

In addition to the bioluminescence measurements, SDS-PAGE analysis of SC-LgBiT and ST-SmBiT mixture confirmed dimer formation mediated by SpyTag-SpyCatcher interaction (**Fig. 2d**). Expectedly, in the control reaction where SC-LgBiT was mixed with SmBiT, no evidence of dimer formation was found (**Fig. S3**). Interestingly, we observed monomeric ST-SmBiT and SC-LgBiT in our SDS-PAGE analysis indicating that only a portion of ST-SmBiT and SC-LgBiT molecules formed heterodimers. This observation is aligned with our previous work^41^ on SpyTag-SpyCatcher-mediated sfCherry reconstitution in which we noted inefficient dimer formation as well. Additionally, our evidence on inefficient NanoBiT reconstitution provided a possible reason for the lower bioluminescence of NanoBiT compared to NLuc (**Fig. S1c**). Nevertheless, since reconstituted NanoBiT demonstrated bright bioluminescence and complete dependence on SpyTag-SpyCatcher interaction for its functional reconstitution, we selected ST-SmBiT and SC-LgBiT as building blocks of our contact-dependent signal generation scheme.

### SspB-iLID dimerization by external illumination in synthetic cells

Once we determined the signal-generation elements in our signaling cascade, we sought to assess the functionality of our signal reception molecule, iLID. We started by asking whether membrane-bound iLID inside synthetic cells can be activated by external blue light illumination. In the presence of cytosolic SspB-mCherry, we expected to observe SspB-mCherry recruitment to the synthetic cell membrane through SspB-iLID dimerization upon blue light irradiation (**Fig. 3a**).

**Figure 3.**
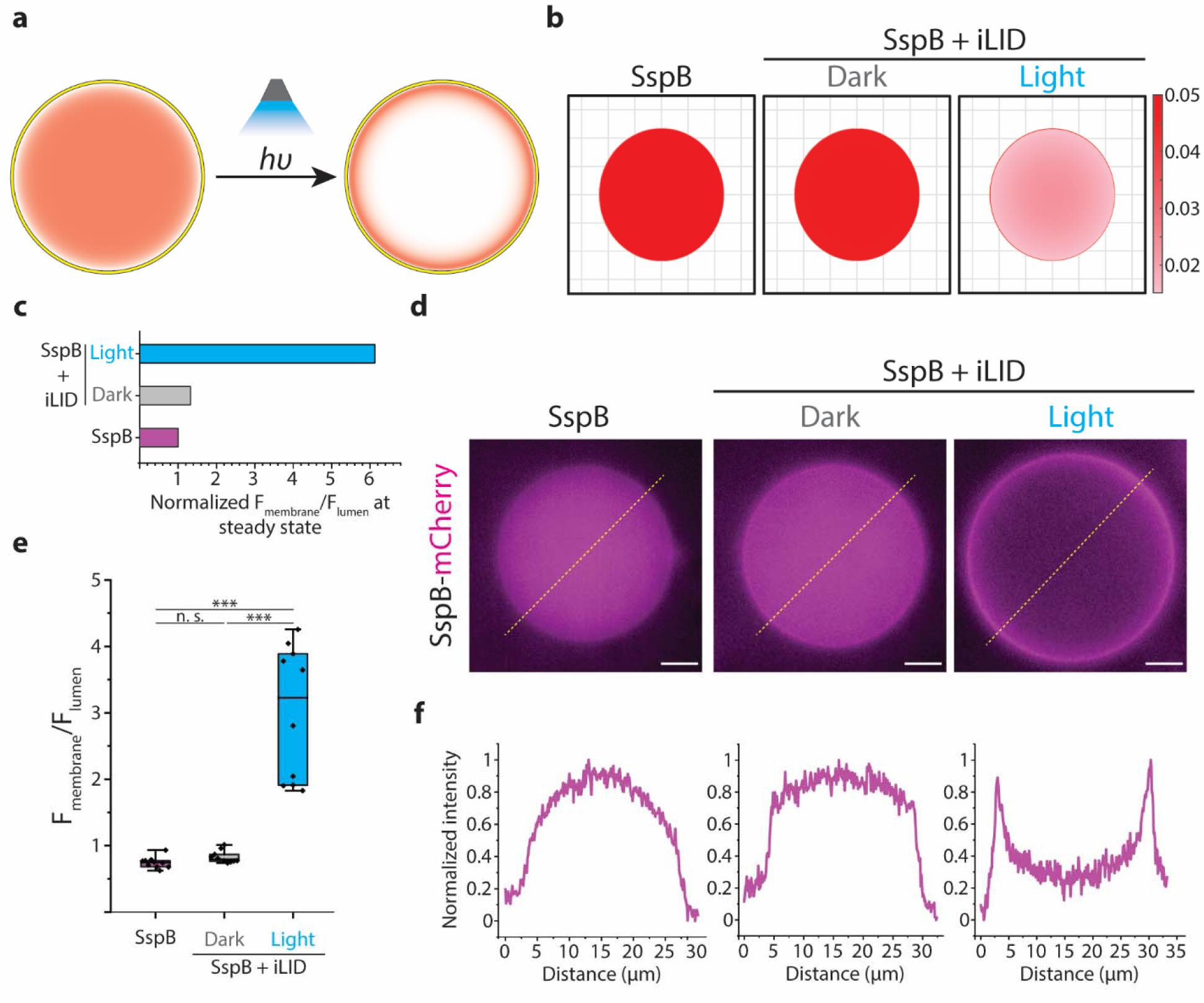
Light illumination drives SspB and iLID interaction inside a GUV. **a**, Schematic representing iLID and SspB encapsulated inside a GUV. SspB-iLID dimerization is induced by external blue light. SspB is illustrated with red color in the lumen of GUV. The depiction of membrane-bound iLID is omitted for simplicity. **b,** Representative results of the mathematical model calculating steady-state SspB concentration inside a 20 μm GUV either encapsulated alone or co-encapsulated with membrane-bound iLID and exposed to dark or light. **c,** Bar graph depicting the normalized ratio of SspB-iLID to luminal SspB at the steady state calculated based on the mathematical model. **d,** Representative fluorescence images of SspB-mCherry encapsulated in GUV (left), SspB-mCherry co-encapsulated in GUV with membrane-bound iLID in the dark (middle), and SspB-mCherry co-encapsulated in GUV with membrane-bound iLID and exposed to external blue light (right). Scale bars: 5 μm. **e,** Boxplots comparing the ratio of SspB-mCherry fluorescence intensity at the GUV membrane to the luminal fluorescence intensity in GUVs encapsulating SspB-mCherry, co-encapsulating SspB-mCherry and iLID in the dark, or co-encapsulating SspB-mCherry and iLID exposed to blue light. The data shows the average ratio of SspB-mCherry signal at the membrane of the GUV to the luminal SspB-mCherry signal with background subtraction for 30 points for ten different GUVs. The box represents the 25–75th percentiles, and the median is indicated. The whiskers show the minimum and maximum data points, *n* = 3. **f,** Intensity profiles of the SspB-mCherry along the dashed yellow line in **d**. Each plot corresponds to the image above in **d**. *p*-values are calculated using one-way ANOVA test and corrected using Tukey’s Honest Significant Difference. n.s. denotes not significant and *** represents *p* < 0.001.

We used a theoretical approach to evaluate whether the amount of SspB-mCherry recruitment to the synthetic cell membrane is enough such that it allows detection of SspB-mCherry at the membrane using fluorescence microscopy. We simulated SspB-mCherry membrane translocation by modeling the SspB-iLID dimerization reaction as well as SspB-mCherry diffusion inside synthetic cells. Based on the work of Guntas *et al*. on iLID development, we chose association constants for iLID-SspB dimerization that reflected their high affinity under the blue light illumination (see Methods section)^45^. Using a finite difference method, we simulated SspB-iLID dimerization both in dark and under blue light stimulation and computed the steady-state location of SspB-mCherry inside a 20 μm diameter circle (**Fig. 3b**).

Our modeling clearly showed that SspB-mCherry recruitment to the membrane occurs only in the presence of both iLID and light illumination (**Fig. 3b**, right). Additionally, the ratio of SspB molecules at the membrane to cytosolic SspB molecules in the presence of light was estimated to be around 6 (**Fig. 3c**). This ratio would allow us to visualize SspB-mCherry recruitment to the membrane via fluorescence microscopy. To test our hypothesis, we generated synthetic cells with phospholipid membranes made of POPC and 5% DGS-NTA (Ni). POPC vesicles were reported to have superior light transmission properties compared to other phospholipid molecules that are commonly used for synthetic cell generation^35^.

We encapsulated 150 nM iLID and 450 nM SspB-mCherry inside synthetic cells and used fluorescence microscopy to detect SspB-mCherry. While SspB-mCherry remained in the lumen of the synthetic cell, iLID was recruited to the membrane due to its C-terminal polyhistidine-tag affinity to NTA (Ni) group on DGS-NTA (Ni) lipids in synthetic cell membrane. Next, we exposed the synthetic cells to external blue light illumination for 15 min before detecting SspB-mCherry. Indeed, we observed strong membrane translocation of SspB-mCherry only in external blue light-exposed synthetic cells that encapsulated membrane-bound iLID (**Fig. 3d**).

Our quantification of SspB-mCherry inside synthetic cells showed that the mCherry intensity at the membrane of synthetic cells encapsulating iLID and exposed to blue light was on average 3 times higher than luminal SspB-mCherry intensity (**Fig. 3e**). This ratio is half of what our model predicted, and the discrepancy can be attributed to the 3D effects that are present only in experimental system as well as possible inefficiencies of iLID activation in experiments that are absent in modeling.

Importantly, iLID-SspB-mCherry dimerization revealed itself in the fluorescence intensity profile of SspB-mCherry along the diameter of the synthetic cell as clear peaks in signal intensity at the membrane that is present only for stimulated synthetic cells encapsulating both iLID and SspB-mCherry (**Fig. 3f**, right). A sudden increase in signal intensity along the synthetic cell diameter can also be noticed in non-stimulated synthetic cells containing iLID and SspB-mCherry which suggests some iLID-SspB-mCherry dimerization in the dark (**Fig. 3f**, middle). On the other hand, synthetic cells encapsulating only SspB-mCherry did not exhibit a peak nor such a sudden increase in their fluorescence intensity profile analysis (**Fig. 3f**, left). Taken together, our results indicate that in our synthetic cell system, membrane-bound iLID can be photoactivated by external blue light as demonstrated through cytosolic SspB-mCherry translocation to the synthetic cell membrane.

### NLuc-mediated SspB-iLID dimerization

While iLID-SspB dimerization was successfully driven by external light illumination, in our designed contact-dependent signal transduction scheme, iLID excitation was required to occur through a bioluminescence signal from a reconstituted luciferase (**Fig. 1c**). Therefore, we set out to test if iLID can be activated by bioluminescence in synthetic cells. To generate bioluminescence, we used NLuc as the luciferase due to its bright and sustained signal. Photoactivation of LOV domain by various luciferases including NLuc has been shown both in natural and synthetic cells. While Kim *et al.* demonstrated that LOV activation by NLuc through bioluminescence resonance energy transfer required NLuc and LOV proximity on the same side of the membrane (*cis* activation)^46^, Chakraborty *et al*. reported that encapsulated RLuc could activate iLID even when the two molecules reside on different sides of the membrane (*trans*-activation)^32^. Since the signal from luciferase must cross the membrane to activate the iLID in our design, we next tested if NLuc can successfully induce SspB-mCherry membrane translocation when it is either inside or outside of the synthetic cell.

First, we purified NLuc with a C-terminal polyhistidine tag and confirmed its peak bioluminescence around 450 nm (**Fig. S4**) which is compatible with iLID excitation wavelength^45^. Next, we encapsulated NLuc, iLID, and SspB-mCherry in synthetic cells with membranes composed of POPC and DGS-NTA (Ni), allowing binding of NLuc and iLID to the membrane due to the polyhistidine tag and NTA (Ni) affinity. We hypothesized that the close proximity between NLuc and iLID on the membrane would lead to photoactivation of iLID by NLuc bioluminescence and subsequent SspB-mCherry membrane translocation (**Fig. 4a**). Consistent with our expectation, we observed that upon addition of furimazine, which is a membrane permeable substrate, SspB-mCherry was recruited to the membrane (**Fig. 4b**). Interestingly, the extent of SspB-mCherry membrane translocation was lower when iLID was excited by intracellular NLuc (**Fig. 4c**) compared to its excitation by external light (**Fig. 3e**).

**Figure 4.**
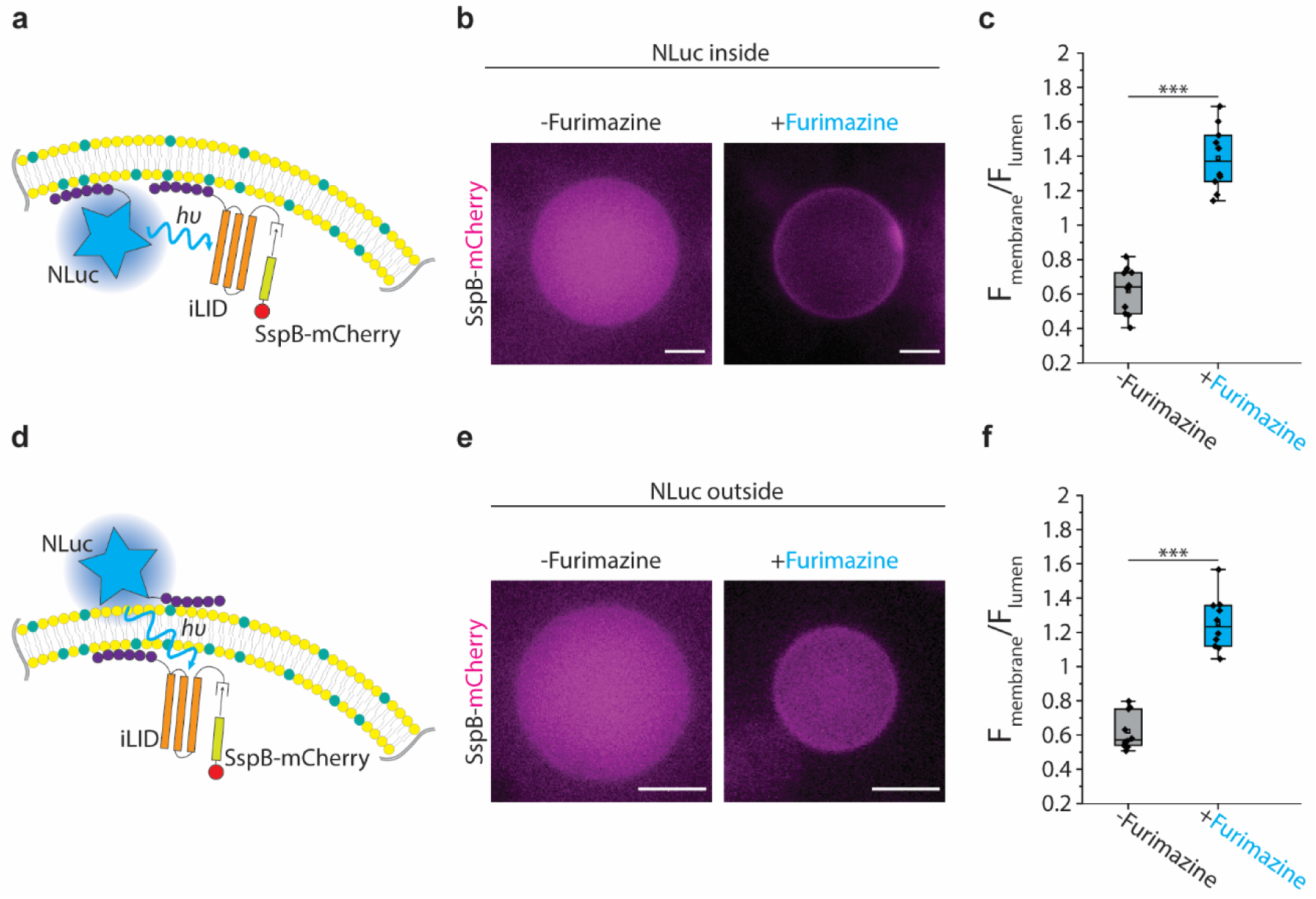
Membrane-bound NLuc drives iLID and SspB interaction. Schematic illustrating membrane-bound iLID activation by NLuc linked to the inner **(a)** or outer **(d)** membrane of GUV that drives SspB-mCherry dimerization with iLID. Representative fluorescence images of SspB-mCherry co-encapsulated with iLID and NLuc in a GUV with NLuc inside **(b)** or outside **(e)** in the absence (left) or presence (right) of NLuc substrate, furimazine. Scale bars: 5 μm. Boxplots comparing the ratio of SspB-mCherry fluorescence intensity at the membrane of GUV to the luminal SspB-mCherry fluorescence intensity with NLuc inside **(c)** or outside **(f)** in the absence or presence of furimazine. The data shows the average ratio of SspB-mCherry signal at the membrane of the GUV to the luminal SspB-mCherry signal with background subtraction for 30 points for ten different GUVs. The box represents the 25–75th percentiles, and the median is indicated. The whiskers show the minimum and maximum data points, *n* = 3. *p*-values are calculated using two-tailed Welch’s *t*-test. *** represents *p* < 0.001.

Upon confirming iLID *cis*-activation, we sought to see whether *trans*-activation of iLID by NLuc is possible in our synthetic cell system (**Fig. 4d**) because iLID excitation by NanoBiT occurs through *trans*-activation in our signal transduction design. We encapsulated iLID and SspB-mCherry in POPC synthetic cells doped with DGS-NTA (Ni) and mixed the synthetic cells with 500 nM of His-tagged NLuc and incubated the mixture to ensure binding of NLuc to the synthetic cell outer membrane. Then, we washed the synthetic cell mixture to remove unbound NLuc from the outer solution. Upon addition of furimazine, we observed SspB-mCherry recruitment to the membrane (**Fig. 4e**), indicating *trans*-activation of iLID by NLuc linked to the outer membrane of a synthetic cell. Our control experiments confirmed that the SspB-mCherry translocation was indeed induced by the membrane-bound NLuc as exclusion of either NLuc or NTA (Ni) lipids caused no SspB-mCherry membrane recruitment (**Figs. S5a** and **S5b**). We note that similar to when NLuc was encapsulated inside synthetic cells, the extent of NLuc-mediated iLID activation was found to be significantly lower than activation by external light (**Figs. 4f** and **S6**). The weaker activation of iLID by NLuc in comparison with external light could be caused by both the lower number of photons emitted by NLuc as well as the constant decay of NLuc photons due to substrate exhaustion.

Further, bioluminescence measurements of NLuc bound to either inside or outside of synthetic cells revealed different kinetics of NLuc bioluminescence (**Fig. S7**). We measured an amplified signal from NLuc linked to the outer membrane of synthetic cells compared to encapsulated NLuc. On the other hand, encapsulated NLuc demonstrated prolonged signal generation. The difference in the amplitude and kinetics of bioluminescence was due to the fact that 500 nM NLuc in the synthetic cell inner solution contains fewer NLuc molecules compared to the outer solution. On the other hand, the concentration of added substrate in both cases was the same. Therefore, the prolonged bioluminescence of encapsulated NLuc was caused by excessive substrate present in the outer solution. Taken together, our results demonstrate the ability of NLuc to activate iLID by both *cis-* and *trans*-activation mechanisms. Thus, we hypothesized that replacing NLuc with NanoBiT would similarly lead to iLID activation, thereby realizing our designed contact-dependent light-based signaling mechanism.

### Contact-dependent light-based synthetic cell communication

Our characterization of individual modules for signal generation and reception allows the demonstration of contact-dependent light-based communication among synthetic cells. To do so, we generated two distinct populations of synthetic cells made by POPC and a small amount of DGS-NTA (Ni). Sender cells encapsulated FITC, a green fluorescence dye, for identification purposes and harbored SC-LgBiT on their outer membrane. Receiver cells, on the other hand, were decorated with ST-SmBiT on their outer membrane while encapsulating membrane-bound iLID and luminal SspB-mCherry.

While SpyTag-SpyCatcher interaction brings the NanoBiT fragments close and promotes formation of the functional luciferase in solution, we found that it was unable to adhere neighboring sending and receiver cells for detectable SspB-mCherry membrane recruitment in the presence of furimazine. To circumvent this challenge, we resorted to using electrostatic attraction between lipid headgroups mediated by salts that has been demonstrated to cause adhesion between lipid membranes^47,48^. We rationalized that in a hypertonic solution of NaCl, the adhesion between deflated synthetic cells will form large interfaces with ample NanoBiT reconstitution, thereby allowing visualization of SspB-mCherry membrane translocation (**Fig. 5a**).

**Figure 5.**
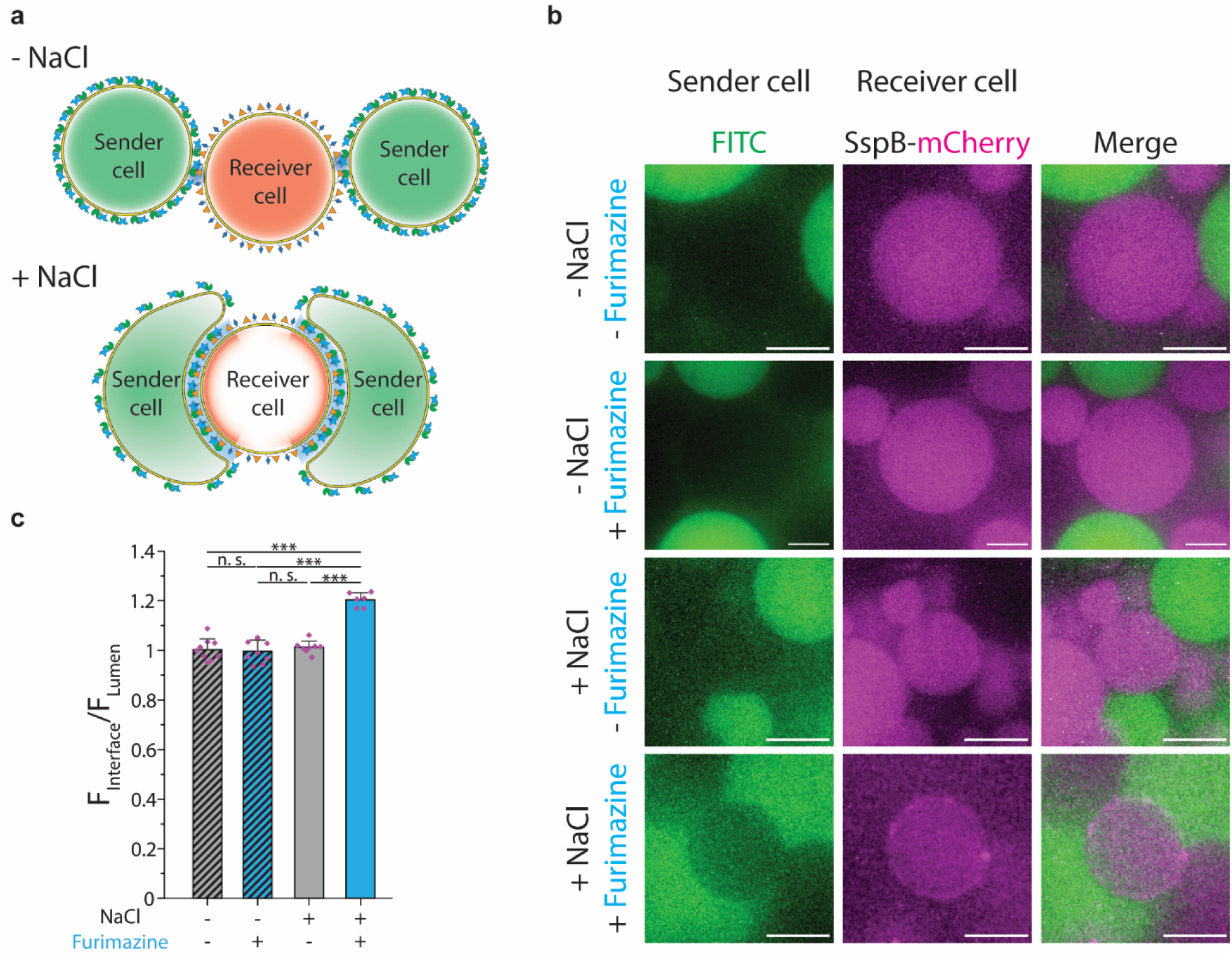
Contact-dependent light-based communication between synthetic cells. **a**, Schematic of light-based contact-dependent communication between synthetic cells. Top: in the absence of NaCl, the small contact area between sender and receiver cells depicted in green and red, respectively, does not result in sufficient SpyTag-SpyCatcher-mediated NanoBiT reconstitution to induce detectable dimerization of SspB-mCherry with iLID. Bottom: in the presence of NaCl, deflated GUVs form large interfaces that cause sufficient SpyTag-SpyCatcher-assisted NanoBiT reconstitution. Functional reconstitution of NanoBiT then activates membrane-bound iLID inside receiver cells that leads to SspB-mCherry recruitment to the receiver cell inner membrane. SspB-mCherry is illustrated in red and the depiction of iLID is omitted for simplicity. **b,** Representative confocal images of FITC (green) encapsulated in sender cells and SspB-mCherry (magenta) encapsulated in receiver cells. iLID activation and SspB-mCherry membrane localization only occur in the presence of both NaCl and furimazine. Scale bars: 10 μm. **c,** Bar plots comparing the ratio of the SspB-mCherry fluorescence intensity at the GUV membrane over the luminal SspB-mCherry fluorescence intensity in the presence or absence of NaCl and furimazine. The data shows the average ratio of SspB-mCherry signal to the luminal SspB-mCherry signal for membrane-membrane interfaces of sender and receiver cells for at least three different receiver cells. The error bars show the standard deviation, *n* = 3. *p*-values are calculated using two-way ANOVA test and corrected using Tukey’s Honest Significant Difference. n.s. denotes not significant and *** represents *p* < 0.001.

We mixed equal amounts of sender and receiver cells in the presence or absence of both NaCl and furimazine and probed whether SspB-mCherry was recruited to the membrane. Confocal fluorescence microscopy images of receiver synthetic cells revealed SspB-mCherry membrane translocation only in the case where furimazine was added to the mixture of synthetic cells in the presence of NaCl (**Fig. 5b**). Analyzing the SspB-mCherry fluorescence along the interfaces of synthetic cells further verified the significant increase in the amount of SspB-mCherry at the membrane of receiver cells that were exposed to both NaCl and furimazine (**Fig. 5c**). In contrast, our control experiment with receiver cells harboring SmBiT, lacking SpyTag domain, did not lead to any change in SspB-mCherry localization, confirming that the SspB-mCherry membrane translocation is due to the NanoBiT reconstitution mediated by SpyTag-SpyCatcher interaction (**Fig. S8a** and **S8b**).

## Discussion

Contact-dependent signaling and communication evolved to enable precise targeting and highly specific communication between cells, making essential processes such as differentiation and phagocytosis possible. With the increasing interest towards synthetic multicellular systems, development of a synthetic contact-dependent communication tool promises biomimicry of various biologically relevant functionalities in synthetic tissues that unlocks synthetic cell applications in regenerative medicine and drug delivery. We demonstrated the design and implementation of a protein-based light-activated contact-dependent communication system that enables contact-dependent signaling between synthetic cells.

In order to make the signaling dependent on physical contact between synthetic cells, we selected a split luciferase protein, NanoBiT, as a signal-generating module and made its function contingent on protein-protein interaction. We leveraged SpyTag-SpyCatcher interaction to permanently bring together the two fragments of NanoBiT and demonstrated SpyTag-SpyCatcher-mediated functional reconstitution of NanoBiT. Inside receiver cells, we encapsulated a light-switchable protein, iLID, and showed its activation triggered by signals coming either from an external light source or a luciferase protein. Lastly, we combined signal generation and reception modules and demonstrated NanoBiT reconstitution in the sender and receiver synthetic cell interfaces which led to a molecular translocation event in receiver cells.

Limited by the simplicity of GUVs as our model synthetic cells, we relied on salt-mediated electrostatic attraction between lipid headgroups to adhere synthetic cells to each other. An avenue worth exploring in the future to improve our design is decorating the GUV membrane with adhesion molecules such as E-cadherin or claudin ^49^ to mimic cell adhesion in synthetic cell communities. In addition, we reported that compared to excitation by external light, NLuc- or NanoBiT-mediated iLID activation was attenuated (**Fig S6**). This is possibly linked to inefficient energy transfer through radiative and non-radiative means and incomplete NanoBiT reconstitution (**Figs. 2d** and **S3**). To circumvent inefficient signaling, one can imagine signaling amplification schemes in which a light-sensitive enzyme, such as a light-sensitive kinase^50^, is activated by light and continues the signaling cascade by phosphorylation of downstream targets.

Previously, we engineered a protein-based tool called InterSpy for the reconstitution of a split fluorescent protein at the membrane-membrane interface between natural and synthetic cells. Here, we extended InterSpy to accommodate signaling between synthetic cells. We chose light as the signal as it does not require auxiliary proteins such as channels and transporters to cross the lipid membrane. Further, the expansive repertoire of optogenetics proteins allows modular designs of the signaling cascade, thus widening the applications. In practice, our designed signaling cascade is analogous to an AND gate which requires physical contact and furimazine to generate light as its output.

While we showed light activation of iLID and SspB membrane recruitment as the response to signal in receiving cells, more complicated responses can be engineered using this AND gate by switching iLID with appropriate optogenetics proteins. For instance, using light-activated transcription elements such as light-sensitive T7^51^ or bacterial light-inducible transcription factor EL222^52^, one can recapitulate contact-dependent induction of gene expression, a process that resembles Notch-Delta signaling, the practical demonstration of which awaits further studies. Lipid-modifying enzymes, such as phosphoinositide (PI) kinases or phosphatases, can be recruited to the contact site to regulate PI synthesis and degradation which is crucial in cellular signal transduction in natural biological systems^53^. Contact-dependent signal generation can also be used to reconstruct a “synthetic synapse” in the future by coupling it with a light-gated ion channel such as channelrhodopsin to allow electrical excitation of cells in the presence of furimazine.

A foundational element of our signaling design was utilizing NanoBiT for signal generation. A split protein that requires protein-protein interaction for its function allowed us to make light generation conditional on physical contact between synthetic cells. Following this strategy, orthogonal signaling cascades can be engineered by replacing NanoBiT with different enzymes that have split variants. For example, replacing NanoBiT with a split TEV protease allows for contact-dependent chemical signaling in which a messenger molecule outside cells can enter cells only if it is processed by the reconstituted split TEV protease^54^. Finally, the recent development of light-activated SpyLigation^55^ which allows SpyTag-SpyCatcher bond formation only in the presence of light can make the activation of orthogonal signaling cascades dependent on each other, thus paving the way for designing signaling schemes with feedback and implementation of protein-based circuits in contact interfaces between natural and synthetic cells.

## Methods

### Cloning and preparation of DNA constructs

Plasmids containing the sequences of iLID and SspB Nano-mCherry were kindly gifted by Dr. Kristen Verhey (University of Michigan). Plasmids with DNA sequences of LgBiT and NLuc were generous gifts from Dr. Taekjip Ha (Harvard Medical School) and Dr. Gary Luker (University of Michigan), respectively. An MBP-SUMO vector was a gift from Dr. Christopher Lima (Sloan Kettering Institute). Target sequences were amplified using Q5 high-fidelity DNA polymerase (NEB). All primers are specified in **Table S1**. Amplified DNA fragments were assembled using Gibson Assembly.

First, the sequences encoding for LgBiT and SpyCatcher003 were amplified from the original LgBiT plasmid and sCatch-GFP plasmid (Addgene #186902^41^), respectively, and were cloned into a pET28b vector to create SpyCatcher-LgBit-6xHis (SC-LgBiT) construct. To generate the SpyTag-SmBiT construct, first, the MBP sequence from MBP-SUMO vector with an N-terminal 6xHis tag was amplified. A g-Block encoding for SmBiT-SpyTag with flexible linkers between SmBiT and SpyTag and at the N-terminus of the SmBiT was ordered from Integrated DNA Technologies (IDT). Next, the g-Block and the amplified MBP were cloned into the pET28b vector to generate 6xHis-MBP-SmBiT-SpyTag (ST-SmBiT) construct. The control construct SmBiT lacking the SpyTag domain was generated by a two-step PCR where a C-terminal SmBiT domain was added to the MBP sequence before cloning into the pET28b vector. Similarly, the iLID construct was generated by cloning amplified MBP-SUMO and iLID sequences and cloning them into pET28b vector to create MBP-SUMO-iLID-6xHis. The amplified SspB-mCherry fragment was cloned into a pGEX-6P-1 vector (from our previous studies^56,57^) to generate GST-mCherry-SspB. Lastly, the NLuc and MBP sequences were amplified and cloned into pET28b vector to generate NLuc-MBP-6xHis.

All cloning sequences were verified by Sanger sequencing (Eurofins). The assembled DNA constructs were purified from XL10-Gold ultracompetent cells (Agilent) using miniprep kits (Qiagen).

### Protein expression and purification

SC-LgBiT and iLID were purified following the conventional His-purification protocol reported elsewhere^41^. Plasmids encoding for the SC-LgBiT or iLID were transformed into BL21-DE3-pLysS competent cells. Single colonies were picked and grown in 5 mL LB broth supplemented with 50 μg/mL kanamycin overnight at 37 °C shaking at 220 rpm. Next, the culture was diluted in 1 L LB supplied with 0.8% w/v glucose and 50 μg/mL kanamycin and was grown at 37 °C shaking at 220 rpm until A_600_ reached 0.5-0.6. The culture was then induced with 0.42 mM isopropyl β-D-1-thiogalactopyranoside (IPTG) and incubated at 30 °C with constant shaking at 200 rpm for 4-5 h. The cells were next harvested by centrifugation at 5000 g for 10 min at 4 °C and then resuspended in 30 mL lysis buffer containing 50 mM Tris-HCl (pH 7.4), 300 mM NaCl, 30 mM Imidazole, and 1 mM 4-(2-aminoethyl) benzenesulfonyl fluoride hydrochloride (AEBSF). The cells were then lysed by a sonicator (Branson Sonifier 450) and the lysate was centrifuged at 30000 g for 25 min at 4 °C. The supernatant was then run through an equilibrated 1 mL HisTrap column (Cytiva) on an AKTA start fast protein liquid chromatography (FPLC) system. Next, the column was washed with 15 column volume washing buffer containing 50 mM Tris-HCl (pH 7.4), 300 mM NaCl, and 50 mM Imidazole. The protein was then eluted by 10 column volumes of elution buffer composed of 50 mM Tris-HCl (pH 7.4), 300 mM NaCl, and 300 mM Imidazole and was collected in 1 mL fractions. The quality of protein purification in each fraction was assessed by SDS-PAGE analysis and the fractions with high concentrations of the protein were pooled and dialyzed against 1 L PBS overnight at 4 °C. The protein concentration was measured using NanoDrop (Thermo Fisher Scientific) and the protein was aliquoted and stored at -80 °C until use.

NLuc, ST-SmBiT, and SmBiT were purified following the same steps described above with the following changes. After harvesting, the cells were resuspended in 30 mL lysis buffer containing 20 mM Tris-HCl (pH 7.4), 300 mM NaCl, 1 mM EDTA, and 1 mM AEBSF and then were lysed by sonication. The lysate was centrifuged at 30000 g for 25 min at 4 °C, and the supernatant was passed through an equilibrated 1 mL MBPTrap (Cytiva) on an AKTA start FPLC system. The bound protein was washed with 15 column volumes of washing buffer containing 20 mM Tris-HCl (pH 7.4) and 300 mM NaCl. The protein was then eluted with 10 column volumes of elution buffer composed of 20 mM Tris-HCl (pH 7.4), 300 mM NaCl, and 10 mM maltose. The purification quality was assessed by SDS-PAGE and the fractions with high concentrations of the protein were pooled and dialyzed against 1 L PBS overnight at 4 °C. The protein concentration was next measured using NanoDrop and the protein was aliquoted and stored at -80 ℃ until use.

Lastly, SspB-mCherry was purified by following conventional GST-purification protocols described elsewhere^56,57^. A single colony of transformed BL21-DE3-pLysS competent cells was picked and grown in 5 mL LB supplemented with 100 μg/mL ampicillin overnight at 37 °C shaking at 220 rpm. Next, the culture was diluted in 1 L LB supplied with 100 μg/mL ampicillin and the cells were grown at 37 °C shaking at 220 rpm until A_600_ reached 0.5-0.6. The culture was then induced with 0.1 mM IPTG and incubated at 30 °C with constant shaking at 200 rpm for 4-5 h. The cells were next pelleted by centrifugation at 5000 g for 10 min at 4 °C and then resuspended in 30 mL lysis buffer composed of PBS and 1% Triton-X100, 1 mM AEBSF, and 1 mM DTT. The cells were lysed by sonication, and the cell lysate was centrifuged at 30,000 g for 25 min at 4 °C and the supernatant was loaded onto an equilibrated 1 mL GSTrap (Cytiva) on an AKTA start FPLC system. The column was then washed with 15 column volumes of washing buffer containing PBS and 1 mM DTT, and the bound protein was eluted by 10 column volumes of elution buffer composed of 50 mM Tris-HCl (pH 7.4), 20 mM glutathione, and 1 mM DTT. The purification quality was analyzed with SDS-PAGE and fractions with high concentration of protein were pooled and dialyzed against 1 L PBS overnight at 4 °C. The protein concentration was measured using NanoDrop and the protein was aliquoted and stored at -80 °C until use.

### Bioluminescence measurement

Bioluminescence measurements presented in **Fig. 2c** were performed using a Synergy H1 (BioTek) multimode plate-reader. All measurements were done using 1 s integration time and a gain of 130. For plots in **Figs. 2b** and **c**, 8 μL mixture of different combinations of ST-SmBiT, SC-LgBiT, and SmBiT were prepared for a final concentration of 500 nM. The mixtures incubated for 30 min at room temperature (RT) before being transferred to a 96 well v-shaped bottom plate. Next, 2 μL of 20-fold diluted furimazine stock (Promega) in live cell substrate (LCS) dilution buffer (Nano-Glo^®^ assay, Promega) was added to each well. The plates were shaken for 10 s inside the plate reader and the measurement was performed right after adding the substrate to the solutions. Photon flux measurements presented in **Fig. S1** were extracted from bioluminescence imaging data captured with an IVIS Lumina Series III (Perkin Elmer, Waltham, MA) and analyzed with Living Image 4.5.2. The integration time was set for 1 s with a binning factor of 2 and f-stop value of 2. 50 μL solutions with different concentrations of NLuc and equimolar concentrations of ST-SmBiT and SC-LgBiT were transferred to a 96-well clear flat bottom plate and incubated for 30 min in RT. Next, 12.5 μL 20-fold diluted furimazine stock in LCS buffer was added to each well and the plates were shaken manually before the time series of bioluminescence images were captured every minute for 30 min.

### SDS-PAGE and in-gel imaging

For SDS-PAGE analyses presented in **Figs. 2d** and **S3**, 10 μL solutions of 1 μM ST-SmBiT, SC-LgBiT, and a 1 μM equimolar mixture of ST-SmBiT and SC-LgBiT were prepared and incubated for 30 min at RT. The solutions were then mixed with 3.3 μL of 4x Laemmli buffer containing 10% 2-mercaptoethanol and were incubated at 95 °C for 10 min. Then, the SDS-PAGE gel was run in a 4–20% Bis-Tris polyacrylamide precast gel (Sigma Aldrich). The gel was stained using SimplyBlue stain (Invitrogen) and imaged by a Sapphire Biomolecular Imager (Azure biosystems) with 658/710 nm excitation/emission wavelengths.

### Synthetic cell preparation

In all experiments, GUVs were generated following the protocol described by Eaglesfield *et al*.^58^ with slight modifications detailed in the following. Appropriate amounts of lipids 1-palmitoyl-2-oleoyl-glycero-3-phosphocholine (POPC) and 1,2-dioleoyl-sn-glycero-3-[(N-(5-amino-1-carboxypentyl) iminodiacetic acid) succinyl] (nickel salt) (DGS-NTA (Ni)) (Avanti Polar Lipids) in chloroform were transferred to a glass vial for a final concentration of 500 μM lipid in oil with 95% POPC and 5% DGS-NTA (Ni) composition. The chloroform was then evaporated under a gentle stream of argon. Next, an appropriate amount of light mineral oil (Sigma-Aldrich, Cat#: M5904) was added to the lipid film and vortexed thoroughly to ensure the lipid is dissolved. The lipid-in-oil solution was incubated at 50 °C for 20 min before being vortexed again. Next, 300 μL of the lipid-in-oil solution was gently layered on top of 400 μL outer solution composed of 50 mM Tris-HCl (pH 7.4) and 300 mM glucose in a 1.5 mL microcentrifuge tube, and the oil-water interface was incubated for 1 h at RT. In a separate 1.5 mL microcentrifuge tube, the inner solution (details in the following sections) was mixed with 600 μL lipid-in-oil solution and the solution was pipetted up and down for 1 min to make a uniform emulsion. The emulsion was then added gently on top of the oil layer, and the tube containing the emulsion and the oil-water interface was centrifuged at 2500 g for 10 min at RT. Next, the top 900 μL of lipid-in-oil solution was removed by aspiration followed by the aspiration of the outer solution until the remaining outer solution volume was around 100 μL. The GUVs were then resuspended in the remaining outer solution by gently pipetting up and down.

### iLID-SspB dimerization by external illumination

GUVs were made by encapsulating 20 μL inner solution containing 50 mM Tris-HCl (pH 7.4), 300 mM glucose, 450 nM iLID-6xHis, 100 nM SspB-mCherry, and 10% v/v OptiPrep (Sigma-Aldrich). After collection, 50 μL of GUV solution was transferred to a 96-well clear flat bottom plate and kept in the dark for 30 min before illumination and epifluorescence imaging using a Nikon TiE inverted microscope equipped with an oil immersion Plane Apochromat 60× objective (NA 1.40), a sCMOS camera (Flash 4.0; Hamamatsu Photonics, Japan), and an HBO 100 W/2 mercury bulb. Next, GUVs were illuminated with an excitation wavelength of 488 nm and intensity of *ca.* 70 mW/cm^2^ for 15 min before single images of SspB-mCherry were taken at an excitation wavelength of 561 nm using ImageJ image acquisition software (NIH).

### iLID-SspB dimerization by NLuc

For experiments with NLuc inside synthetic cells, GUVs were generated by encapsulating 20 μL inner solution containing 50 mM Tris-HCl (pH 7.4), 300 mM glucose, 450 nM iLID-6xHis, 100 nM SspB-mCherry, 500 nM NLuc, and 10% v/v OptiPrep. After GUVs were collected, 80 μL of GUV solution was transferred to a 96-well clear flat bottom plate and incubated in the dark for 30 min. Next, 20 μL LCS buffer containing a 20-fold dilution of furimazine stock was added to the well and epifluorescence images of SspB-mCherry were taken immediately. A 100-fold dilution of furimazine stock was added to the well every 15 min for a total of three times during imaging.

For experiments with NLuc attached to the outer membrane of synthetic cells, GUVs were made by encapsulating 20 μL inner solution containing 50 mM Tris-HCl (pH 7.4), 300 mM glucose, 450 nM iLID-6xHis, 100 nM SspB-mCherry, and 10% v/v OptiPrep. After GUVs were collected, 500 nM NLuc was added to the GUV solution and GUVs were incubated at RT for 30 min. Next, 900 μL outer solution was added to the GUV solution and the solution was centrifuged at 2500 g for 10 min at RT. Next, 900 μL of outer solution was removed by gentle pipetting. The GUV pellet was then resuspended by gently pipetting up and down and 80 μL of GUV solution was transferred to a 96 well clear flat bottom plate. Then, 20 μL LCS buffer containing a 20-fold dilution of furimazine stock was added to the well and epifluorescence images of SspB-mCherry were taken immediately. A 100-fold dilution of furimazine stock was added to the well every 15 min for a total of three times during imaging.

### iLID-SspB dimerization by NanoBiT

Sender cells were generated by encapsulating 20 μL inner solution containing 50 mM Tris-HCl (pH 7.4), 500 mM KCl, 12.5 μM Fluorescein isothiocyanate (FITC)-dextran, and 10% v/v OptiPrep in an outer solution containing 50 mM Tris-HCl (pH 7.4) and 1 M glucose. Similarly, receiver cells were made by encapsulating 20 μL inner solution containing 50 mM Tris-HCl (pH 7.4), 500 mM KCl, 450 nM iLID-6xHis, 100 nM SspB-mCherry, and 10% v/v OptiPrep in an outer solution containing 50 mM Tris-HCl (pH 7.4) and 1 M glucose. After collecting GUVs, 5 μM SC-LgBiT and ST-SmBiT were added to sender and receiver cells, respectively, in individual 1.5 mL microcentrifuge tubes and GUV solutions were incubated at RT for 30 min. Next, the GUVs were washed by adding 900 μL outer solution to each tube followed by centrifugation at 2500 g for 10 min at RT. Then, 900 μL of supernatant from each tube was removed gently by pipetting. Each population of GUVs was then resuspended and a 100 μL 1:1 mixture of sender cells and receiver cells was made and transferred to a 96-well clear flat bottom plate. Next, 100 μL 1 M NaCl was added to the well and the GUVs were incubated for 30 min in the dark to let membrane-membrane interfaces and NanoBiT form. GUV images were taken using an oil immersion Plan-Apochromat 60 x/1.4 NA (Olympus) objective on an inverted microscope (Olympus IX-81) equipped with an iXON3 EMCCD camera (Andor Technology), National Instrument DAQ-MX controlled laser (Solamere Technology), and a Yokogawa CSU-X1 spinning disk confocal unit. Images were acquired using MetaMorph (Molecular Devices). Single images of FITC and SspB-mCherry were taken at excitation wavelengths of 488 and 561 nm, respectively. Then, 50 μL LCS buffer containing a 20-fold dilution of furimazine stock was added to the well, and images of FITC and SspB-mCherry were taken immediately. A 100-fold dilution of furimazine stock was added to the well every 15 min during imaging.

### Image analysis

All GUV images were analyzed by ImageJ. The intensity profiles presented in **Fig. 3f** were obtained by measuring the fluorescence intensity along the corresponding indicated lines in **Fig. 3d**. In addition, the background signal was measured and subtracted from the intensity values followed by signal intensity normalization to the maximum signal. For analyzing membrane recruitment of SspB-mCherry presented in **Figs. 3e**, **4c**, **4f**, and **S5**, ImageJ Oval_profile plugin was used to measure the mCherry fluorescence intensity over 30 points equally spaced along the periphery of the GUV. Similarly, the fluorescence intensity of GUV lumen and background were measured and averaged. For measuring SspB-mCherry recruitment to GUV-GUV interfaces presented in **Figs. 5c** and **S8b**, the fluorescence intensity along 5 arbitrary lines crossing the GUV-GUV interfaces were measured and the average signal of three pixels centered around the peak intensity and the average signal of its 10 following points into the lumen of the receiver cell were calculated as the signal values associated with membrane and lumen, respectively. When there was no clear peak in the intensity profile (as in control cases), the average of 3 intensities centered around the point where the signal reached a plateau and the average signal of 10 following points into the lumen of a receiver cell were taken as membrane and lumen intensity values, respectively. All 5 measurements were then averaged and represented a single data point.

### Mathematical modeling

A 1-dimensional mathematical model was developed to model the interaction of SspB-mCherry with membrane-bound iLID inside a GUV with radius R under the effect of radial diffusion and chemical reaction between iLID and SspB. For simplicity, the angular diffusion was neglected. The diffusion along the radius of the GUV was mathematically modeled with the following equation:

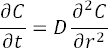

Where *C*, *r*, *D*, and *t* represent the concentration of SspB-mCherry, the radius, the diffusion coefficient, and time, respectively. The equation requires an initial condition and two boundary conditions. A Neumann boundary condition at the *r* = 0 was assumed:

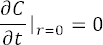

This boundary condition implies that the flow of diffusion will always be from the center of the GUV towards the membrane. At the membrane (*r* = R), the boundary condition is coupled to the chemical reaction between SspB and iLID. If the concentration of iLID and SspB-iLID dimer is represented by *L* and *M*, respectively, the chemical reaction can be modeled with the following set of equations:

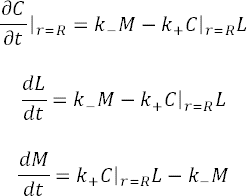

in which, *k_+_* and *k_-_* represent the association and dissociation rate constants of iLID and SspB dimerization. For simplicity, it was assumed that under light illumination, only *k_+_* changes. However, the ratio of *k_+_* to *k_-_* follows experimental data reported by Guntas *et al*.^45^ That is, *k_+_* in light is 36 times higher than *k_+_* in the dark. Since the above partial differential equation (PDE) has a boundary condition in the form of a set of ordinary differential equations (ODE), it cannot be solved analytically. Therefore, a finite difference model was developed to solve the PDE and ODEs simultaneously using Matlab. With small timesteps, it was reasoned that in each step, the ODE can be solved first, thus generating a fixed boundary condition for PDE in the next step, and this cycle can be repeated until the solution is convergent.

### Finite-difference modeling of ODEs

Assuming u_0_, u_1_, and u_2_ represent the concertation of SspB, iLID, and SspB-iLID dimer at the membrane, respectively, at time *t*, after a small timestep Δt, the new concentrations can be calculated by the following equations:

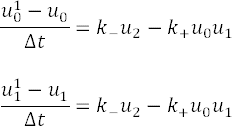

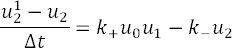

in which, the variables with superscript 1 represent the value of the variable at time *t* + Δt.

### Finite-difference modeling of PDEs

The diffusion PDE can be reduced to algebraic equations using a finite-difference approach assuming that the GUV radius is divided into N equal elements. If *C_n_* represents the concentration of SspB in the n^th^ element, the PDE will be reduced to the following equation:

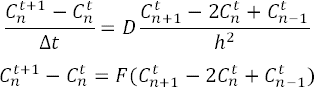

where

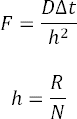

and t+1 and t superscripts stand for the concentration of SspB at the time *t* + Δt and *t*, respectively. At the first element, we can apply the Neumann boundary condition which gives us the following:

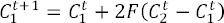

To apply the Neumann boundary condition at *r* = R, the rate of change in concentration of SspB at this element is considered to be constant and can be calculated from the set of ODEs at *r* = R:

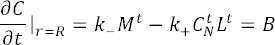

where C_N_ stands for the concentration of SspB at the N^th^ element. Applying this boundary condition to the finite-difference model results in the following:

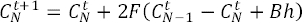

If all C_n_ at time *t* and *t* + Δt are represented in column vectors **C^t^** and **C^t+1^**, respectively, then the **C^t+1^** can be found from the relationship between **C^t^** and **C^t+1^**:

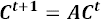

in which **A** is the following N by N matrix:

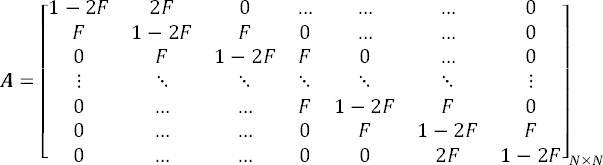

In each step, after finding the value of **C^t+1^**, 2FBh is added to the c^t+1^ to account for the boundary condition. This approach was implemented in a Matlab code to solve the diffusion PDE coupled with boundary condition ODEs with the initial condition of 0.05, 1.5, and 0 for the concentration of SspB in each element, iLID at the N^th^ element, and SspB-iLID dimer at N^th^ element, respectively. The results presented in **Fig. 3b** were plotted using Matlab. The bar graph in **Fig. 3c** was made by obtaining the final values of the solution for luminal SspB and SspB-iLID dimer.

### Statistical analysis

Experiments were performed in at least three independent replicates. Bioluminescence data presented in **Fig. 2c** was transferred to Microsoft Excel and Welch’s *t*-test was done to compare the maximum bioluminescence between the control and experiment group using the Excel’s built-in data analysis tool. The *p-*values reported in **Fig. S1c** were calculated by Welch’s t-test as well. The statistical significance between the ratio of membrane-bound SspB-mCherry to luminal SspB-mCherry in the presence and absence of furimazine presented in **Figs. 4c** and **f** and **S5b** was also calculated by Welch’s *t*-test. In addition, an R code was written and used to perform one-way and two-way ANOVA tests on data from analyzed images represented in **Figs. 3d** and **5b**, respectively, with Tukey’s Honest Significance Difference correction. Confidence intervals of regression lines presented in **Fig. S2b** and **S2c** were calculated using OriginPro 2020b statistical analysis tool. All *p*-values are listed in **Table S2**.

## Acknowledgment

We thank Dr. Tobias Pirzer (Technical University of Munich), Dr. Christopher Lima (Sloan Kettering Institute), Dr. Taekjjp Ha (Harvard Medical School), Dr. Gary Luker (University of Michigan), and Dr. Kristin Verhey (University of Michigan) for plasmids (see **Table S1**). We thank Dr. Gary Luker (University of Michigan) and Dr. Kenneth Ho (University of Michigan) for helping with bioluminescence imaging. The work is funded by the National Science Foundation (EF-1935265) and the National Institutes of Health (R01-EB030031).

## Notes

### Competing Interest Statement

The authors have declared no competing interest.

