## Supplemental information for "Light-based juxtacrine signaling between synthetic cells"

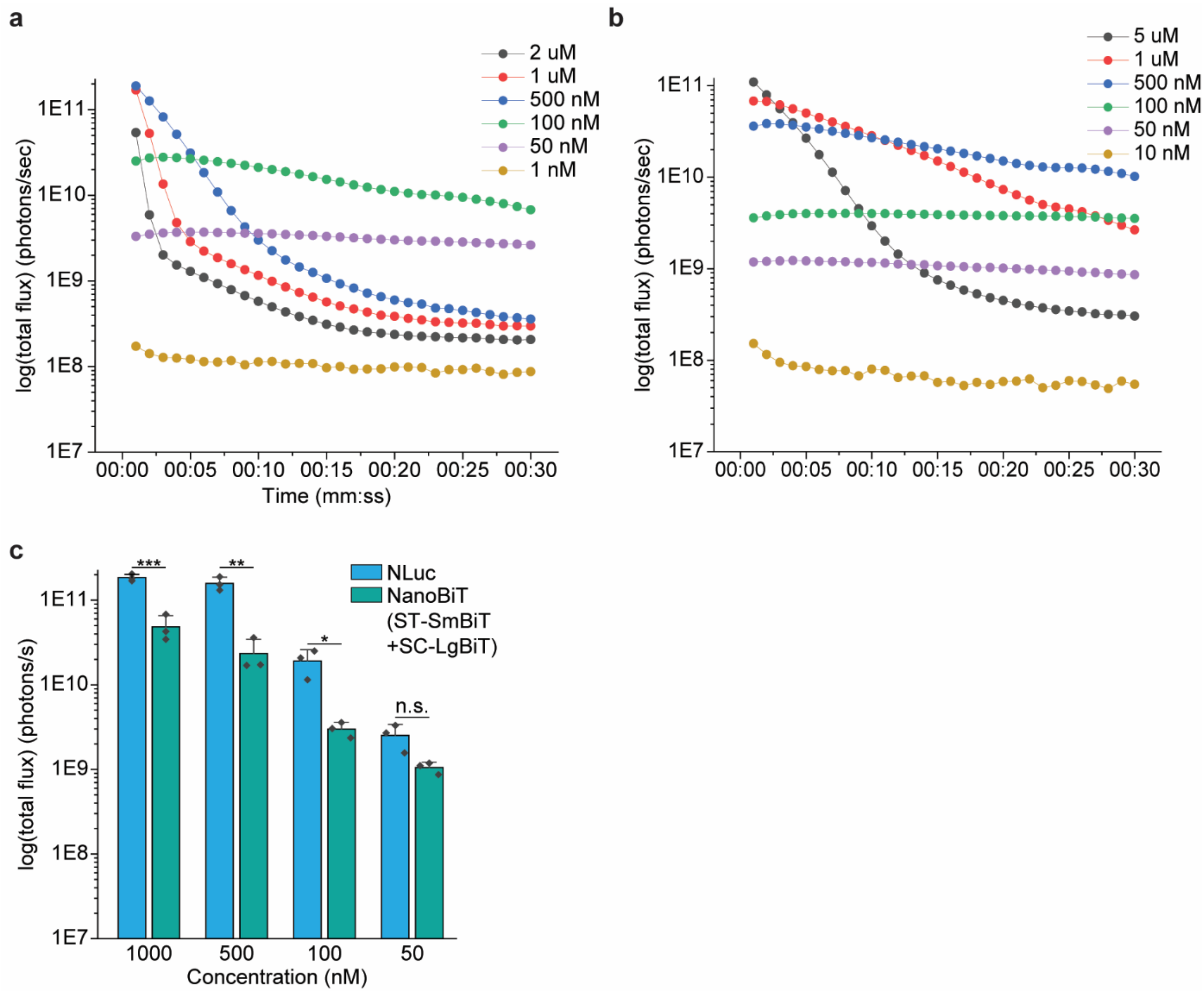


**Figure S1| Bioluminescence imaging of NLuc and reconstituted NanoBiT reactions.** Representative time series depicting quantification of bioluminescence imaging of 50 µL reactions with different NLuc **(a)** or NanoBiT **(b)** concentrations in the presence of 1:4 dilution of live cell substrate (LCS) buffer containing 20-fold dilution of furimazine stock. **c,** Bar plots comparing the maximum bioluminescence at different concentrations of NLuc and NanoBiT. Error bars show the standard deviation. *p*-values are calculated using one-tailed Welch's t-test, $n = 3$. ***, **, *, and n.s. represent $p < 0.001$, $p < 0.01$, $p < 0.05$, and not significant, respectively.

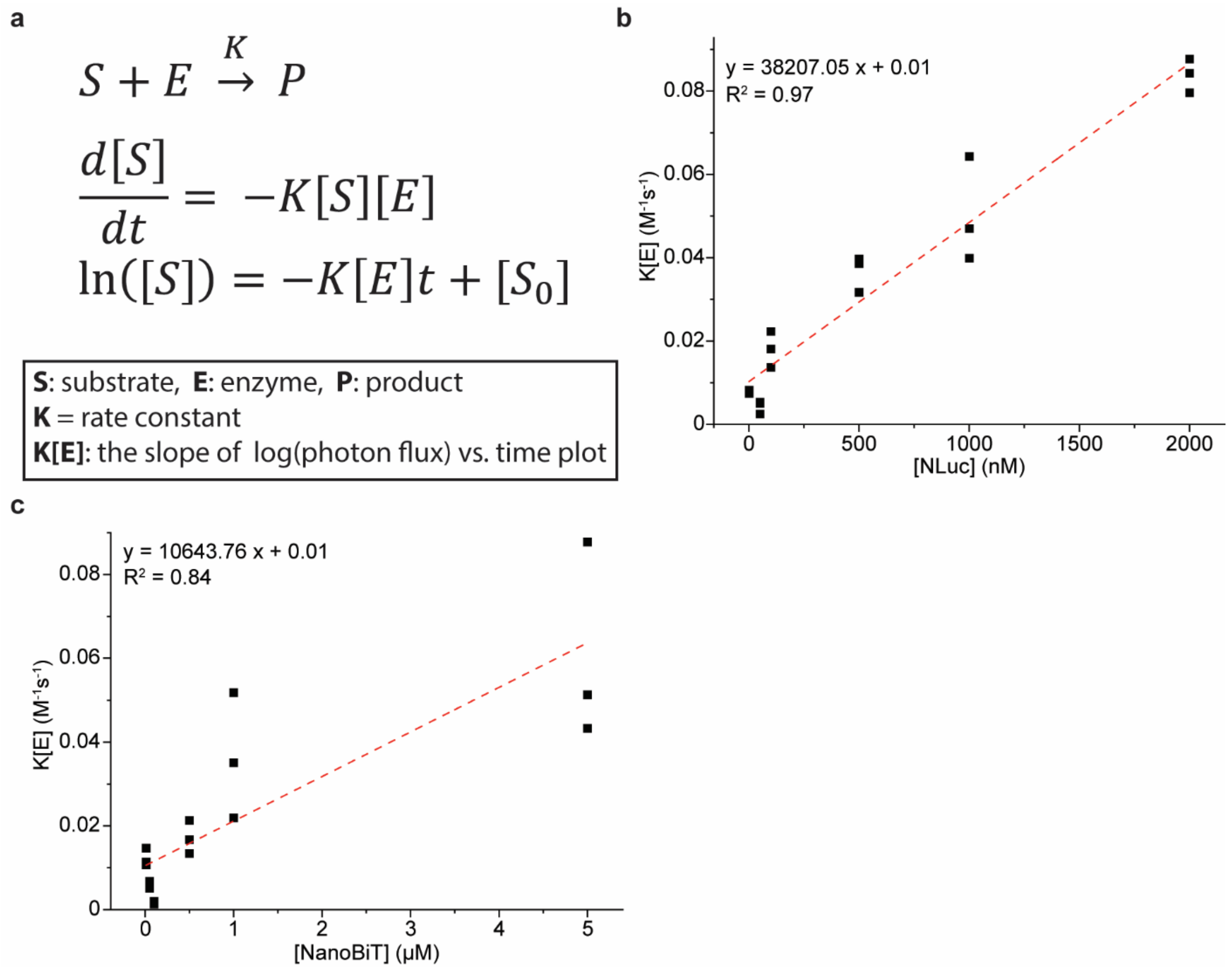


**Figure S2| Numerical modeling of the bioluminescence reaction reveals rate constants of NLuc and NanoBiT. a,** Modeling bioluminescence reaction using differential equations indicates that enzyme rate constant is proportional to the slope of log(photon flux) vs. time plot. Regression analysis on the relationship of the slope of log(photon flux) vs. time plot with NLuc **(b)** or NanoBiT **(c)** concentration determines the enzyme rate constant, $n$ = 3.

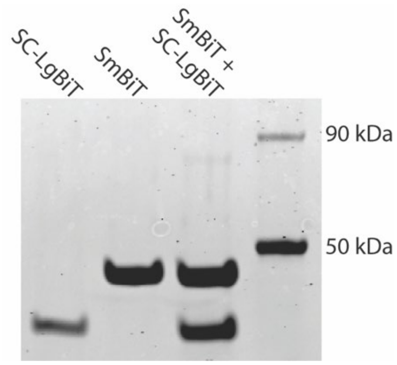


**Figure S3| Absence of SpyTag-SpyCatcher interaction results in no dimer NanoBiT formation.** In-gel fluorescence imaging of the Coomassie-stained SC-LgBiT (lane 1), SmBiT (lane 2), the mixture of SC-LgBiT and SmBiT (lane 3), and the ladder (lane 4).

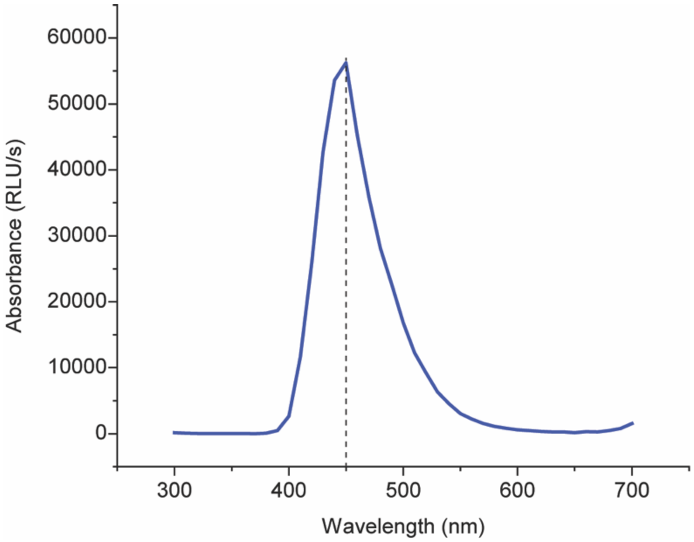


**Figure S4| NLuc bioluminescence peaks at 450 nm.** Bioluminescence emission spectrum of 500 nM NLuc in the presence of 1:4 dilution of LCS buffer containing 20-fold dilution of furimazine stock.

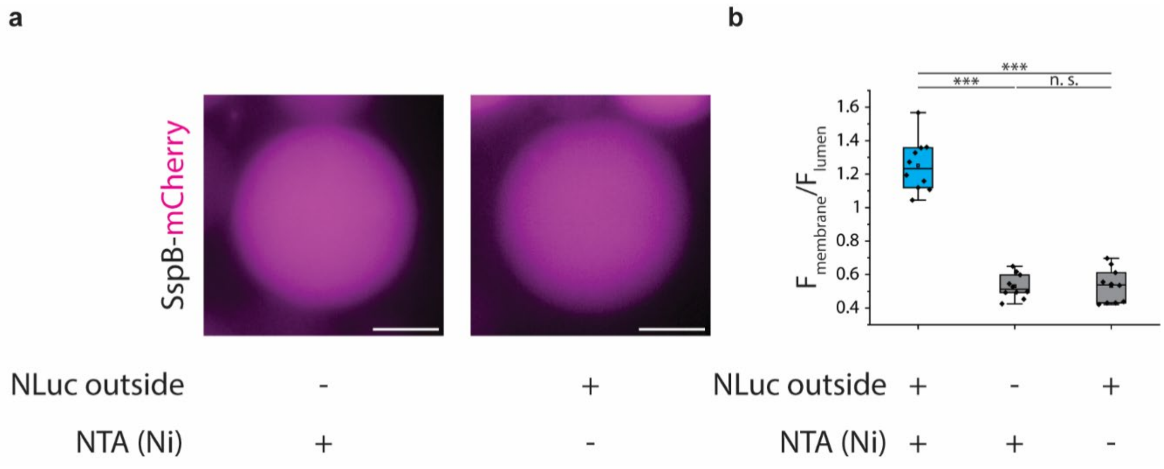


**Figure S5| Absence of NLuc or NTA (Ni) results in no SspB-mCherry membrane recruitment. a,** Representative fluorescence image of a GUV encapsulating SspB-mCherry without addition of NLuc to its outer solution (left) or without including DGS-NTA (Ni) lipid in its membrane composition (right). Scale bar: 10 µm **b,** Box plot comparing the ratio of SspB-mCherry signal at the GUV membrane to the luminal SspB-mCherry signal in POPC and DGS-NTA (Ni) GUVs with or without outer membrane-bound NLuc or POPC GUVs with NLuc. The data shows the average ratio of SspB-mCherry signal at the membrane of the GUV to the luminal SspB-mCherry signal with background subtraction for 30 points for ten different GUVs. The box represents the 25–75th percentiles and the median is indicated. The whiskers show the minimum and maximum data points, $n = 3$. $p$-values are calculated using two-way ANOVA test and corrected using Tukey's Honest Significant Difference. n.s. denotes not significant and *** represents $p < 0.001$.

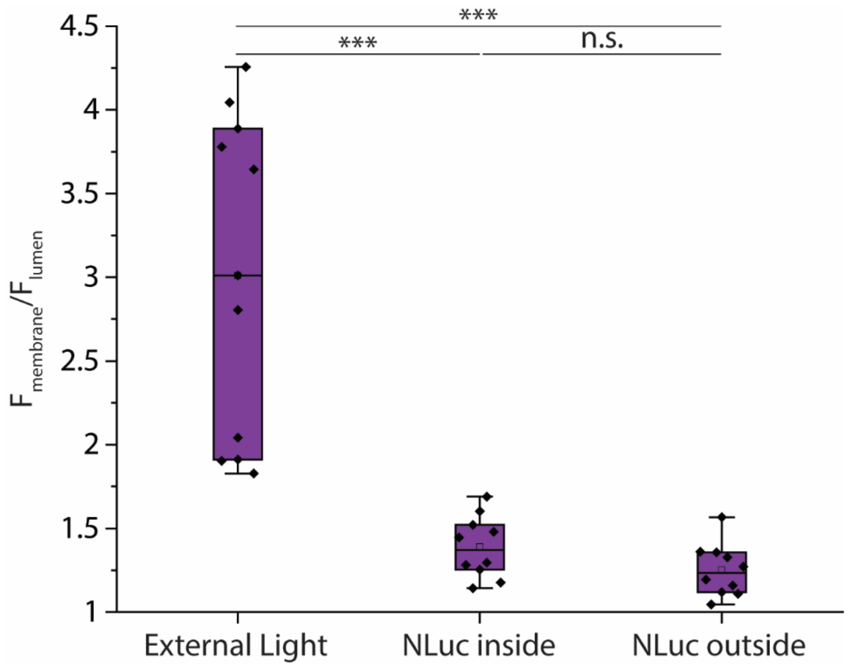


**Figure S6| SspB-mCherry membrane recruitment through iLID activation via an external light source is significantly stronger than NLuc-mediated iLID activation.** Box plot comparing the ratio of SspB-mCherry fluorescence signal at GUV membrane to the luminal SspB-mCherry signal for GUVs encapsulating 450 nM iLID and 100 nM SspB-mCherry excited by an external light source (see Methods) for 15 minutes, 500 nM encapsulated NLuc, or 500 nM NLuc attached to their outer membrane. The data shows the average ratio of SspB-mCherry signal at the membrane of the GUV to the luminal SspB-mCherry signal with background subtraction for 30 points for ten different GUVs. The box represents the 25–75th percentiles, and the median is indicated. The whiskers show the minimum and maximum data points, $n = 3$. $p$-values are calculated using one-way ANOVA test and corrected using Tukey's Honest Significant Difference. n.s. denotes not significant and *** represents $p < 0.001$.

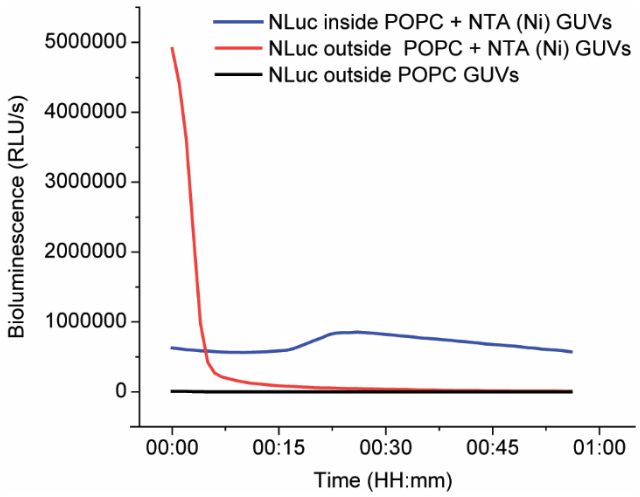


**Figure S7| Encapsulated membrane-bound NLuc demonstrates a different reaction kinetics compared to NLuc attached to the outer membrane of synthetic cell.** Representative bioluminescence readout from 100 µL solutions of GUVs with NLuc either attached to their inner or outer membranes or POPC GUVs washed after incubation with NLuc. Each solution was mixed with 25 µL of LCS buffer containing 20-fold dilution of furimazine stock right before bioluminescence measurement.

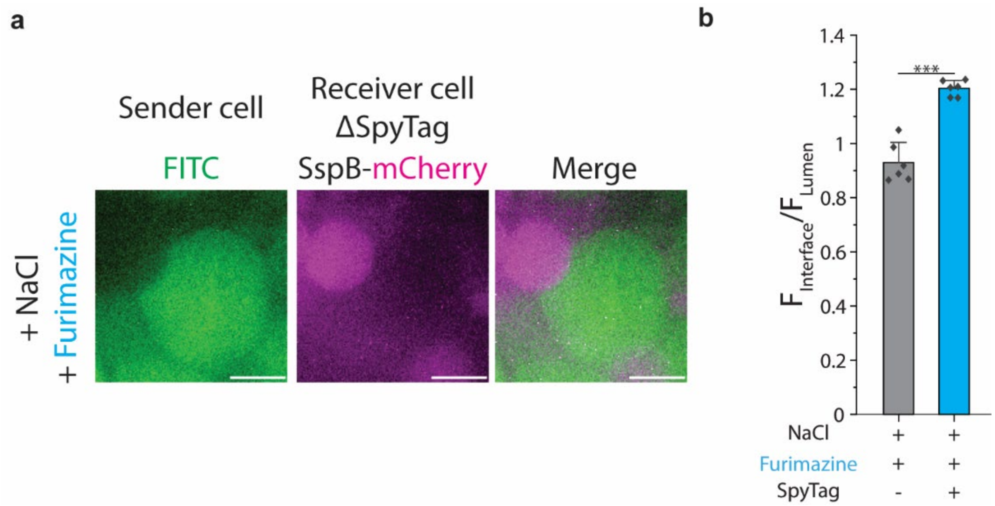


**Figure S8| Deletion of SpyTag domain from ST-SmBiT on receiver cells results in no SspB-mCherry translocation to the membrane of receiver cells. a,** Representative confocal images of FITC (green) encapsulated in a sender cell and SspB-mCherry (magenta) encapsulated in a receiver cell. Scale bars: 10 µm. **b,** Bar plots comparing the ratio of the SspB-mCherry fluorescence intensity at the membrane of a receiver cell to the luminal SspB-mCherry fluorescence intensity in the presence or absence of SpyTag domain in ST-SmBiT. The data shows the average ratio of SspB-mCherry signal to the luminal SspB-mCherry signal for membrane-membrane interfaces of sender and receiver cells for at least three different receiver cells. The error bars show the standard deviation, $n$ = 3. $p$-value is calculated using two-tailed Welch's test. *** represents $p < 0.001$.

**Table S1:** List of all generated constructs and primers used for cloning

| Construct | Fragment | Primers | Template DNA (Source) |
|---|---|---|---|
| pET28b-His-MBP-SmBiT-SpyTag | Vector | **Following primers were used for amplifying vector for all pET28b constructs**<br>**FWD**: CACCACCACCACCACCAC<br>**REV**: AAAAAACCTCCTTACTTTCTAGTCTCAAG | pET28b-RQF (Dr. Tobias Pirzer, Technical University of Munich) |
| | MBP | **FWD**: TCTTGAGACTAGAAAGTAAGGAGGTTTTTTATGCACCACCACCACCACCACggttcaggcATCGAAGAAGGTAAACTGGTAATC<br>**REV**: ATAGCCGGTCACgcctgaaccacctcccgaAGTCTGCGCGTCTTTCAG | pET28b-His-MBP-SUMO (Dr. Christopher Lima, Sloan Kettering Institute) |
| | SmBiT-SpyTag | **IDT duplex oligo**:<br>tcgggaggtggttcaggcGTGACCGGCTATCGCCTGTTTGAAGAAATTCTGgcgggttcaggcggttcaggccgtggcgttcctcatattgttatggtggacgcctacaaacgctataaaTGAGATCCGGCTGCTAACAAAGCCCGAAAG | N/A |
| pET28b-His-MBP-SmBiT | MBP-SmBiT | **Fragment made by 2-step PCR:**<br>**FWD1:**<br>TCTTGAGACTAGAAAGTAAGGAGGTTTTTTATGCACCACCACCACCACCACggttcaggcATCGAAGAAGGTAAACTGGTAATC<br>**REV1:**<br>cgcCAGAATTTCTTCAAACAGGCGATAGCCGGTCACgcctgaaccacctcccgaAGTCTGCGCGTCTTTCAG<br>**FWD2:** TCTTGAGACTAGAAAGTAAGGAGGT<br>**REV2:** TTCCTTTCGGGCTTTGTTAGCAGCCGGATCTCACGCCAGAATTTCTTCAAACAGG | pET28b-His-MBP-SUMO (Dr. Christopher Lima, Sloan Kettering Institute) |
| pET28b-SpyCatch | SpyCatcher-LgBiT | **FWD:** TCTTGAGACTAGAAAGTAAGGAGGTTTTTTATGACCACACTGTCCGGACTG | pcDNA-xC-LgBiT (Dr. Taekjip Ha, |

| | | | |
|---|---|---|---|
| er-LgBiT-His | | **REV:** AGCCGGATCTCAGTGGTGGTGGTGGT GGTGGCTGTTGATGGTTACTCGGAAC | Harvard Medical School) |
| pET28b-iLID-MBP-His | MBP | **FWD**: TCTTGAGACTAGAAAGTAAGGAGG TTTTTTatgATCGAAGAAGGTAAACTGGT **REV**: tgccagtccggatccagagccggagccgccT CCACCAATCTGTTCTCTG | pET28b-His-MBP-SUMO (Dr. Christopher Lima, Sloan Kettering Institute) |
| | iLID | **FWD**: GGTGgaggcggctccggctctggatccgga CTGGCAACCACACTGGAAC **REV**: AGCCGGATCTCAGTGGTGGTGGTG GTGGTGtcccgaaaagtaattttcgtcgttcgctgc | pHR-SFFVp-iLID::EGFP::FTH 1 (Dr. Kristin Verhey, University of Michigan) |
| pET28b-NLuc-MBP-His | MBP | **FWD**: TCTTGAGACTAGAAAGTAAGGAGG TTTTTTATGGTCTTCACACTCGAAGATTTCG **REV**: gcctgaaccacctcccgacgcAGAACCG CTCAGAATCTCCTC | pET28b-His-MBP-SUMO (Dr. Christopher Lima, Sloan Kettering Institute) |
| | NLuc | **FWD**: agcggttctgcgtcgggaggtggttcaggc ATCGAAGAAGGTAAACTGGTAATC **REV**: AGCCGGATCTCAGTGGTGGTGGTG GTGGTGGTGGTGGTGGTGtcccgaAGTCTGC GCGTCTTTCAG | pCMV-CXCL12-NLuc (Dr. Gary Luker, University of Michigan) |
| pGEX-mCherry-SspB | Vector | **FWD**: tagTGACTGACTGACGATCTGCCT **REV**: GGGCCCCTGGAACAGAA | pGEX6p1-Fascin-sfGFP (homemade) |
| | SspB-mCherry | **FWD**: TCGGATCTGGAAGTTCTGTTCCAGG GGCCCATGGTGTCTAAAGGCGAGG **REV**: CGCGCGAGGCAGATCGTCAGTCAGT CActaACCAATATTCAGCTCGTCATAG | pHR-SFFVp-FUSN::mCherry::SspB (Dr. Kristin Verhey, University of Michigan) |

**Tables S2:** List of *p*-values:

| **Figure** | **Comparison groups** | ***p*-value** |
|---|---|---|
| 2-c | ST-SmBiT+SC-LgBiT vs. SmBiT+SC-LgBiT | 0.00015 |
| 3-e | Light vs. Dark | $6.40 \times 10^{-9}$ |
| | Light vs. SspB | $2.90 \times 10^{-9}$ |
| | Dark vs. SspB | 0.94 |
| 4-c | -Furimazine vs. +Furimazine | $7.21 \times 10^{-9}$ |
| 4-f | -Furimazine vs. +Furimazine | $1.49 \times 10^{-8}$ |
| 5-c | -NaCl-Furimazine vs. -NaCl+Furimazine | 0.98 |
| | -NaCl-Furimazine vs. +NaCl+Furimazine | $4.08 \times 10^{-10}$ |
| | +NaCl-Furimazine vs. -NaCl+Furimazine | 0.81 |
| | +NaCl+Furimazine vs. +NaCl-Furimazine | $2.07 \times 10^{-9}$ |
| | +NaCl+Furimazine vs. -NaCl+Furimazine | $2.00 \times 10^{-10}$ |
| S1-c | 1000 nM NLuc vs. 1000 nM NanoBiT | 0.00032 |
| | 500 nM NLuc vs. 500 nM NanoBiT | 0.0024 |
| | 100 nM NLuc vs. 100 nM NanoBiT | 0.028 |
| | 50 nM NLuc vs. 50 nM NanoBiT | 0.052 |
| S5-b | +NLuc+NTA (Ni) vs. -NLuc+NTA (Ni) | $2.32 \times 10^{-13}$ |
| | +NLuc+NTA (Ni) vs. +NLuc-NTA (Ni) | $2.51 \times 10^{-13}$ |
| | +NLuc-NTA (Ni) vs. -NLuc+NTA (Ni) | 0.99 |
| S6 | External light vs. NLuc inside | $2.44 \times 10^{-6}$ |
| | External light vs. NLuc outside | $5.97 \times 10^{-7}$ |
| | NLuc inside vs. NLuc outside | 0.86 |
| S8 | +SpyTag vs. -SpyTag | $6.50 \times 10^{-5}$ |
